# Microbiome and resistome dynamics along a sewage-effluent-reservoir continuum underline the role of natural attenuation in effluent receiving reservoirs

**DOI:** 10.1101/2022.09.21.508967

**Authors:** Inês Leão, Leron Khalifa, Nicolas Gallois, Ivone Vaz-Moreira, Uli Klümper, Daniel Youdkes, Shaked Palmony, Lotan Dagai, Thomas U. Berendonk, Christophe Merlin, Célia M. Manaia, Eddie Cytryn

## Abstract

This study assessed temporal dynamics of total and antibiotic resistant fecal bacterial indicators and antibiotic resistance genes (ARG) along a sewage-effluent-reservoir continuum, in an experimental system consisting of a sewage-fed membrane-aerated bioreactor (MABR) whose effluent fed a 4500 L polypropylene basin that mimicked an effluent storage reservoir. We applied a multidisciplinary approach that coupled physicochemical analyses, cultivation of total and cefotaxime-resistant E. coli, microbiome (bacterial and eukaryotic) analysis and qPCR/ddPCR quantification of selected ARGs. Total and cefotaxime-resistant E. coli loads dropped by approximately 1.5 log units in both the MABR and the reservoir, but the relative reduction (normalized to 16S rRNA genes) in both E. coli and ARGs was higher in the reservoir. Reservoir microbiomes were significantly different from those in the MABR, and were characterized by temporal shifts and periodic algal (Chlorophyta) blooms that were coupled to oxygen and pH fluctuations. Collectively, the data indicates that the decrease in E. coli and ARGs in the MABR was primarily facilitated by sludge removal, whereas in the reservoir, it was predominantly associated with microbial community shifts. The study highlights the capacity of ecological interactions in mitigating antibiotic resistance in both engineered and natural ecosystems.

**Importance:** Antibiotic resistance is considered one of the most significant public health predicaments of the 21st century, and there is growing evidence that anthropogenically impacted environments such as those receiving raw and treated sewage can contribute to this phenomenon. In this study, we evaluated the dynamics of total and antibiotic resistant fecal pathogen indicators and antibiotic resistance genes along a sewage-treated wastewater-effluent reservoir continuum, concurrent to evaluation of microbial community composition and physicochemical parameters. Results indicate that both the treatment bioreactor and the effluent storage reservoir removed resistant bacteria and antibiotic resistance encoding genes. However, in the reactor removal was primarily linked to physical processes, whereas in the storage reservoir it appeared to be strongly facilitated by ecological interactions. The study highlights the capacity of aquatic ecosystems to alleviate antibiotic resistance, and suggests that ecological processes in aquatic ecosystems can be harnessed to mitigate antibiotic resistance.

## Introduction

Raw sewage is a mirror of the gut microbiota of the served population (1), and is consequently a reservoir of fecal-derived antibiotic resistant bacteria (ARB) and resistance genes (ARGs), whose abundance and diversity vary as a function of geography and socioeconomic conditions (2, 3). Although ARGs are minor components of the sewage metagenome (0.03%), the sewage resistome constitutes a vast array of genetic determinants that confer resistance to the entire spectrum of antibiotic classes, including ARGs associated with emerging clinical threats(2, 4). Conventional wastewater treatment processes generally remove 1-3 log-units ml^-1^ (per volume) of fecal bacteria, including ARBs and associated ARGs, and this removal can be augmented by disinfection (5). Nonetheless, wastewater treatment plant (WWTP) effluents frequently contain substantial ARB and ARG loads (6–11). These determinants can potentially disseminate through the water cycle and the food chain, after being discharged to aquatic ecosystems, or following irrigation with reclaimed wastewater, and contribute to clinically associated antibiotic resistance. (2, 12–14).

Fecal bacterial indicators that are routinely targeted for water quality assessment (15) provide no insights regarding antibiotic resistance. This can be overcome by monitoring fecal bacterial indicators resistant to next-generation antibiotics concomitant to total counts (16), as proposed in a recent review (17). Source tracking of ARGs using quantitative PCR-based methods provide a rapid and accurate means of quantification that sheds light on antibiotic resistance levels in WWTPs and receiving environments, but there are close to 3000 documented antibiotic resistance determinants (18), and currently there are no established ARG standards for assessing water quality. Within an epidemiological context, ARGs are only interesting to monitor if they can be horizontally transferred to other bacteria and/or they are associated with pathogens. With this respect, Zhang et al. (19) recently proposed ranking ARGs according to the associated risk level for human health, where rank I encompasses ARGs that are associated with MGEs and are present in ESKAPE pathogens (20). The limitations of both cultivation-based and culture-independent molecular analyses underline the fact that combining the two is imperative for holistic understanding of antibiotic resistance in WWTP effluents and downstream environments.

Various studies have investigated the fate of WWTP effluent-derived ARB and ARG in receiving aquatic ecosystems such as rivers, lakes and effluent stabilization reservoirs used for storing treated wastewater prior to irrigation (11, 20–22). While certain studies indicate that ARGs can persist in receiving water and sediment (21), others suggest that they are either diluted or actively removed (11). It has been proposed that mitigation of sewage derived ARB and ARGs is associated with ecological interactions (22), but the scope and nature of these interactions are still not well understood due to environmental complexity and difficulty of source tracking (20).

The goal of this study was to elucidate the scope, dynamics and potential mechanisms responsible for ARB and ARG removal in a large-scale pilot system containing a semi-commercial scale membrane bioreactor treating municipal sewage, whose secondary effluent fed a large (4500 L) reservoir where attenuation or enhancement of antibiotic resistance may occur. We applied a holistic analytical pipeline that measured physicochemical analyses, cultivation of total and antibiotic resistant fecal coliforms, microbiome (bacterial and eukaryotic) analysis and quantification of potentially hazardous ARGs and associated MGEs markers (qPCR and ddPCR) along the sewage-effluent-reservoir continuum. The closed nature of the system and the coupling of isolation, culture-independent ARG quantification and analysis of microbial community composition provided important insights into the mechanisms potentially responsible for mitigation antibiotic resistance in WWTPs and receiving aquatic environments.

## Materials and Methods

### Description of experimental system

Sampling campaigns were conducted between July and December 2020 on an experimental wastewater treatment beta-site (**Figure. 1**), situated within the Maayn Zvi municipal wastewater treatment plant (32.59684, 34.92975) in Israel. The system consisted of an Aspiral L3 (www.fluencecorp.com/wp-content/uploads/2018/05/Aspiral-Product-Brochure.pdf) Membrane Aerated Biofilm Reactor (MABR) connected to a cylindrical polypropylene reservoir (4500 L working volume), aimed to mimic operational reservoirs commonly used for effluent storage prior to irrigation. The passive aeration by diffusion of oxygen through MABR membranes supports an aerobic nitrifying biofilm that develops on their surface, while suspended solids are held in the mixed liquor, enabling simultaneous nitrification and denitrification. The feed flow rate of primary effluent to the MABR was approximately 5 m^3^/h with slight variations due to occasional equipment failures (*i.e*. clogged feed pump, ruptured diffuser), power failures (4 overall) and excess sludge removal, which reduced the desired effluent quality. Mixing frequency and duration, Return Activated Sludge (RAS), Sludge wasting (WAS) and aeration regimes were modulated to maintain bioreactor performance. Reservoir retention time was initially 21 days (September 23^rd^, 2020 to October 14^th^, 2020), after which it was reduced to 10 days (October 14^th^, 2020 to November4^th^, 2020), and later on 5 days (November 4^th^, 2020 to November 25^th^, 2020).

**Figure 1.**
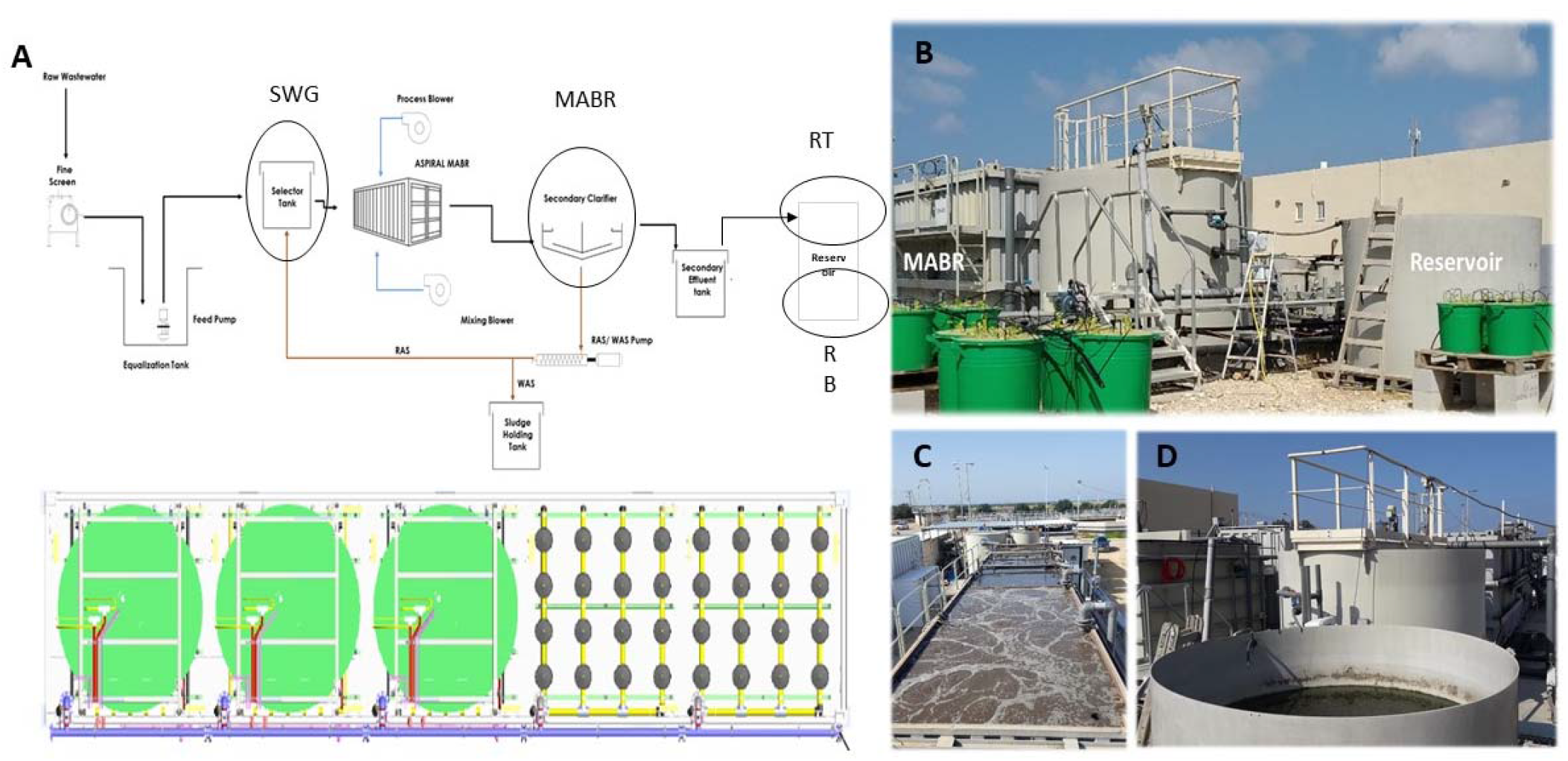
Overview of the experimental beta-site. Schematic diagram of the raw sewage-Membrane Aerated Biofilm Reactor (MABR)-reservoir continuum with sampling points circled in black (**A-top**). Aerial diagram of the MABR (**A-bottom**), where large green circles on the left represent three MABR modules, and the small black circles represent fine-bubble diffusers. Profile (**B**) and aerial (**C** and **D**) photo of the MABR and reservoir at the beta-site. SWG-raw sewage; MABR-membrane aerated bioreactor; RT (reservoir top): Sampling faucet situated at the top 10 cm of the reservoir; RB (reservoir bottom): Sampling faucet situated at the bottom 10 cm of the reservoir.

Samples for physicochemical, bacterial and molecular analyses (**Table S1**) were taken from faucets situated at different points along the sewage-MABR-reservoir continuum (**Figure. 1**). Faucets were opened for 30s and sampling vessels were washed 3 times before sampling to remove residual water in the pipes. Water samples for bacterial enumeration and community DNA extraction were either immediately filtered on site, or transferred on ice to the lab at the Volcani Institute and filtered within 3 h.

### Physical and Chemical analyses

Temperature, oxygen, pH and conductivity were measured using a HQ40D model digital two channel multi meter (HACH, CO, USA) using specific electrodes for each parameter, and turbidity was measured with a 2100Q Portable turbidimeter (HACH, CO, USA). Total organic carbon (TOC) was determined by dry combustion with a Flashea™ 1112 NC elemental analyzer (Thermo Fisher Scientific, Hanau, Germany). Ammonia, nitrite, nitrate, phosphorus and sulfate were measured colorimetrically with a Quickchem 8000 Autoanalyzer (Lachat Instruments, Milwaukee, WI) using standard protocols provided by the manufacturer.

### Microbial quantification, isolation and characterization

Cultivation-based analysis was applied to enumerate total and cefotaxime-resistant coliform and *Escherichia coli* in the raw sewage, MABR effluent and reservoir, using a modified version of the standard membrane filtration method (ISO 9308-1). Briefly, ten-fold serial dilutions (10^-2^ to 10^-5^, and 10^0^ to 10^-4^ for total and cefotaxime resistant coliforms, respectively) of the collected raw sewage and effluent samples were prepared in sterile saline solution (0.85% (w/v) NaCl) and 1 ml of diluted sample was filtered in triplicates through a 0.45 μm nitrocellulose grid membrane filter using a vacuum filtration system. Subsequently, filters were placed (grid facing upwards) on CCA (Coliform Agar acc. to ISO 9308-1 Chromocult^®^) culture media plates with or without cefotaxime (4 mg/L) and plates were incubated at 37 °C overnight. Coliform and *E. coli* colonies on the CCA media were enumerated based on the color classification defined by the supplier.

To validate presumptive CCA colorimetric identifications, 108 randomly-selected colonies from raw sewage, MABR and the reservoir were classified on a microflex LT MALDI-TOF MS system (Bruker Daltonics GmbH, Bremen, Germany) using the Flex Control v3.4 Biotyper automation software as previously described(23). Results supported the manufacturer’s colorimetric taxonomic characterizations, as: 55 blue colonies were identified as *E. coli* or *Shigella* and 53 red colonies were identified as *Klebsiella* spp. and *Enterobacter* spp.

### DNA extraction, storage, and shipment

For DNA extraction, 10 to 100 ml of the freshly collected samples were filtered through 0.22 μm polycarbonate membranes using a filtration unit and then stored at −80 °C prior to extraction. Sampling dates and specific volumes filtered for each sample are shown in **Table S2**. DNA was extracted from these membranes using the DNeasy PowerWater kit (cat# 14900-100-NF, Qiagen, USA), using the protocols provided by the manufacturer. Purified DNA was divided into four aliquots and stored at −80 °C, until shipping. Composite (top and bottom) reservoir samples were prepared by mixing equal volumes of DNA, after top and bottom physicochemical parameters were found to be very similar to each other. Samples for bacterial community analysis and qPCR quantification of ARGs were shipped to Universidade Católica Portuguesa in Porto, Portugal; samples for ddPCR quantification of ARGs and MGEs were shipped to Université de Lorraine, Nancy, France; and samples for eukaryotic community analysis were shipped to Technische Universität Dresden, Germany. All samples were shipped on dry ice by 2-day express delivery and stored at −80 °C upon arrival.

### Quantification of 16S rRNA gene, Mobile Genetic Elements & Antibiotic Resistance Genes

Digital droplet PCR (ddPCR) and quantitative PCR (qPCR) were applied to quantify the bacterial 16S rRNA gene, the *Escherichia coli* indicator gene *uidA* and the CrAssphage fecal contamination indicator, as well as five ARGs (*sul1, ermF, bla*_VIM-2_, *bla*_KPC_, *bla*_CTX-M-1_) and four MGE families (class 1 integrons, *Tn916/Tn1545* ICE family, SXT ICE family, IncP plasmid family). The above-mentioned genes were all analyzed on the samples from the following dates: 19^th^ and 25^th^ of August, 15^th^ of September, 24^th^ of November, 1^st^ and 15^th^ of December 2020. The ddPCR was conducted on a One-Step QX200 system (Biorad Hercules, CA, USA) using the Evagreen or Taqman technologies. The qPCR was performed on a StepOnePlus™ machine (Applied Biosystems, Life Technologies, Carlsbad, CA, USA) using TaqMan technology (**Table S3**). Primers, mix reactions and amplification conditions for ddPCR and qPCR are described in **Table S3**. For ddPCR, the QX Manager™ software (version 1.7.4, BioRad) was used to assign positive/negative droplets after adjusting the thresholds manually, and to convert counts into copies/μL. Negative controls (DNA- and RNA-free water), and a positive control (artificial target DNA) were used in the first ddPCR assay, to confirm the proper position of the positive/negative droplets. Negative controls and a calibration curve were run with each qPCR determination. We initially compared qPCR and ddPCR quantifications of 16S rRNA and *ermF* genes using identical sets of DNA to determine the relation between the two approaches (**Figure S1**).

The rationale for choosing the targeted genes is as follows. The beta-glucuronidase encoding *uidA* gene was chosen because it is frequently used to source track *E. coli* (the most common fecal bacterial indicator) in aquatic ecosystems, and the CrAssphage bacteriophage was selected because it is highly abundant in the human gut. The ARGs were targeted based on expected abundance and ubiquity and different risk levels according to Zhang *et al*. (19), included the widespread *sul1* gene (rank IV), the human-enriched *ermF* gene (rank III/IV), and three ARGs of concern to public health, *bla*_CTX-M-1_ (rank III), *bla*_VIM_-2 (rank I/III), and *bla*_KPC-2_ (rank I). Regarding the rationale for MGE selection, *Tn916/Tn1545* ICE family is abundant in *Bacillota*, the SXT/R391 ICE family is predominant in *Gammaproteobacteria*, IncP-1 conjugative plasmids are profuse in *Pseudomonadota*, and Class 1 integrons are broadly found in Gram-negative bacteria.

### Microbial community analyses

Prokaryotic communities were analyzed by targeting the V3-V4 region of the 16S rRNA gene, using the primers 341F and 806R (**Table S3**). Amplicons were sequenced using an Illumina paired-end platform to generate 250 bp paired-end raw reads that were merged with FLASH (V1.2.7) (24) and quality filtered using QIIME software (V1.7.0). The chimeric sequences were removed and the reads with good quality were assigned to operational taxonomic units (OTUs; ≥97% sequence identity). OTUs annotation was performed against the SSUrRNA SILVA v.138 Database (http://www.arb-silva.de/) (25). The weighted UniFrac distance matrix was used to generate a Principal Coordinate Analysis (PCoA) and a features table used for sample comparison and statistical analysis performed with STAMP software (v2.1.3).

Eukaryotic communities were analyzed by targeting the 18S rRNA gene using the universal primers 1391f and EukBr (26) (**Table S3**) that target the V9 variable region. PCR products with proper size were selected following validation by 2% agarose gel electrophoresis. Equimolar amounts of PCR products from each sample were pooled, end-repaired, A-tailed and further ligated with Illumina adapters. Libraries were sequenced at Novogene Co. (Cambridge, UK) using the Illumina MiSeq platform generating 250 bp paired-end raw reads, which were assigned to samples based on their unique barcodes and truncated by cutting off the barcode and primer sequences. Subsequently, FLASH (24) (v1.2.11, http://ccb.jhu.edu/software/FLASH/) was applied to merge reads, and fastp (27) for quality control of raw tags, to obtain high-quality clean tags. Vsearch software (28) was used to blast clean tags against the database, to detect and remove chimeric sequences. The deblur module in QIIME2 (29) was used to denoise, and sequences with less than 5 reads were filtered out to obtain the final ASVs (Amplicon Sequence Variables) and feature table. Finally, the Classify-sklearn module in QIIME2 software was used to compare ASVs with the SILVA Database (http://www.arb-silva.de/)(25), and to obtain the species annotation of each ASV. In tandem, the weighted UniFrac distance for PCoA analysis was calculated in QIIME2.

The 16S and 18S rRNA sequences were uploaded to the NCBI-SRA archive under Bioproject number PRJNA805207.

### Statistical analyses

One-way ANOVA followed by an LSD post-hoc test was applied to evaluate statistical significance between the raw sewage, MABR and reservoir for each of the measured parameters, and one-way ANOVA followed by a Tukey-Kramer post-hoc test was applied to evaluate temporal variance. For all analyses, differences were considered significant when p-values were below 0.05.

Potential relationships between species-level microbial community composition and structure, ARG&MGE markers, and environmental and physicochemical variables were assessed using Redundancy Analysis with Canoco 5.01 software (30). The relationship between species and environmental variables was assessed based on 1000 Monte Carlo permutations, followed by forward selection with the criterion of p<0.01 of significance.

IBM SPSS Statistics version 28 was used to statistically analyze the alpha diversity of the most abundant prokaryotic (>5%) and eukaryotic microbiomes (>1%) and for comparing the qPCR and ddPCR analyses (UCP and LCPME labs, respectively) of 16S rRNA and *ermF* genes. For alpha diversity, ANOVA was applied with Tukey as post-hoc test with a significance level of p<0.01. For the comparison between laboratory results, a Friedman test was performed (with significance level of p<0.01) because the data did not follow a normal distribution. Statistical calculations for gene abundance and microbial community analyses are shown in **Tables S5-S8**.

## Results

### Physicochemical fluctuations along the sewage-MABR-Reservoir continuum

We evaluated physicochemical parameters along the sewage-MABR-reservoir continuum between July and December 2020, at 17 time points. Water temperatures ranged from 35 to 17 °C. In the November and December profiles, reservoir temperatures were approximately 5 °C lower than the raw sewage (**Figure. 2A**), indicating that they are more strongly impacted by ambient temperatures. The pH (**Figure. 2B**) and dissolved oxygen levels (**Figure. 2C**) were relatively stable in raw sewage and the MABR (except for 22^nd^ July and 8^th^ September 2020 where oxygen in the MABR was low due to system malfunction), but varied more in the reservoir, seemingly due to photosynthetic activity. Turbidity (**Figure. S2A**) was almost completely alleviated in the MABR (with the exception of September 1^st^ and 8^th^, 2020 when malfunctions occurred), correlating to significant reduction in total organic carbon (**Figure. 2D**). Likewise, over 80% of total nitrogen was removed in the MABR (**Figure. 2E**), corresponding to the removal of most of the ammonia (**Figure. 2F**). Mass balance of all the analyzed species indicated that most of the carbon and nitrogen in the system was either gasified (to CO_2_, N_2_ or N_2_O) or removed as settled sludge biomass, considering the fact that nitrate (**Figure. 2H**) and nitrite (**Figure. 2G**) concentrations in the MABR effluent were 1-2 orders of magnitude lower than the influent ammonia concentration. Phosphate levels (**Figure. S2B**) significantly dropped between raw sewage and MABR suggesting biomass accumulation of polyphosphate, and sulfate (**Figure. S2C**) increased between raw sewage and MABR, implying oxidation of reduced sulfur compounds such as H_2_S.

**Figure 2.**
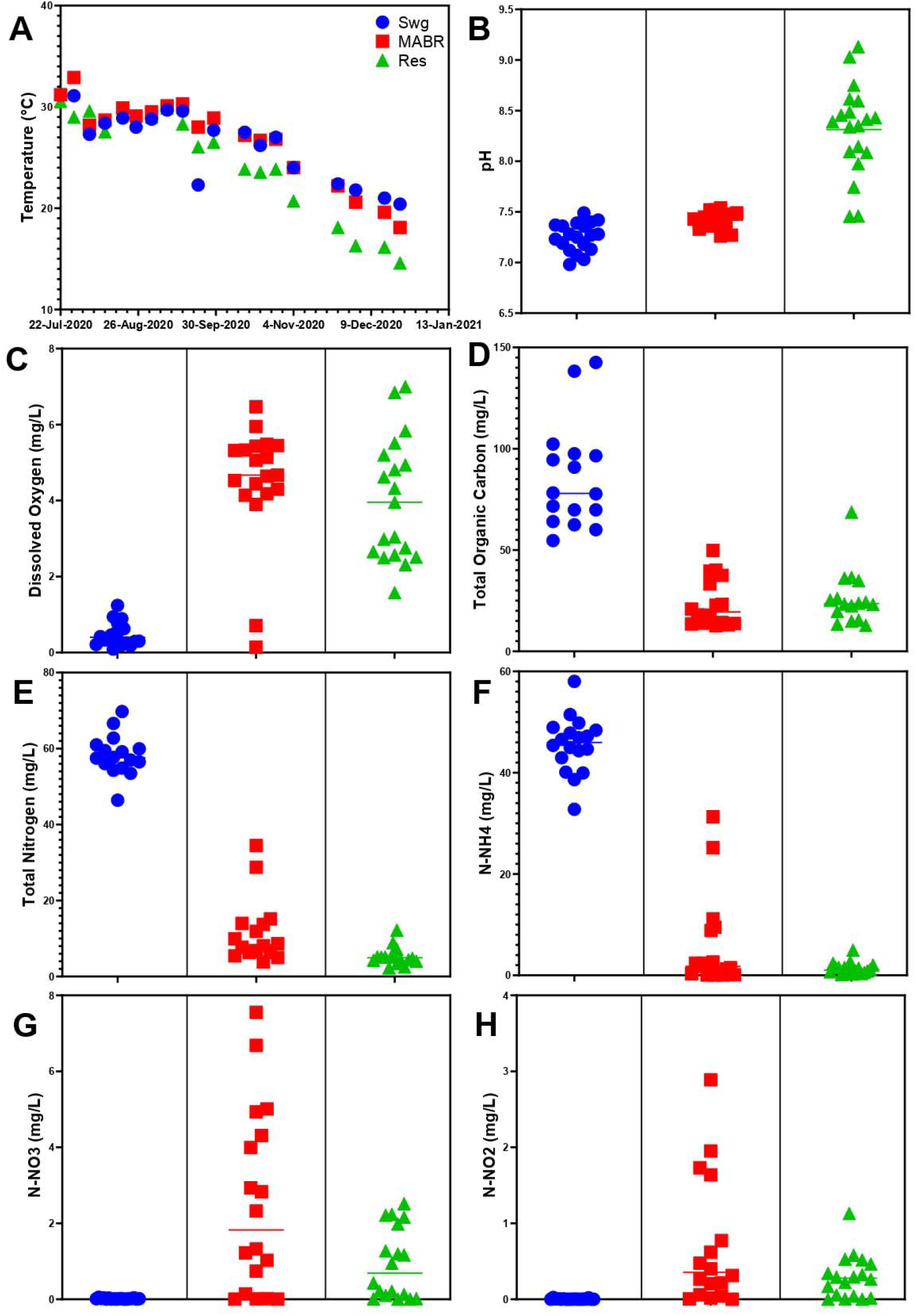
Physicochemical analyses in the raw sewage (blue circles), MABR (red boxes) and reservoir (green triangles). Temperature (A); pH (B); dissolved oxygen (C); total organic carbon (D). Total nitrogen (E); ammonia (F); nitrate (G); nitrite (H).

### Fecal coliform dynamics along the sewage-MABR-Reservoir continuum

We evaluated total and cefotaxime-resistant *E. coli* (**Figure 3A**) and fecal coliforms (**Figure S3**), to determine their levels in raw sewage levels, and their removal in the MABR and reservoir. On average, the abundance of these fecal bacterial indicators decreased by approximately 1.5 log units ml^-1^, although fluctuations in removal capacity were observed at different time points (**Figure S4**). Normalizing to the 16S rRNA gene levels measured by quantitative PCR analyses (**Figure 3B**), revealed that the abundance of *E. coli* relative to the total bacterial community decreased more in the reservoir than in the MABR. *E. coli* values measured in the raw sewage and MABR were similar to levels of the *E. coli* marker gene *uid*A (see below), supporting the culture-based analyses. In contrast, in the reservoir *uid*A levels were slightly higher than cultivated *E. coli* levels, suggesting the presence of non-viable bacteria.

**Figure 3.**
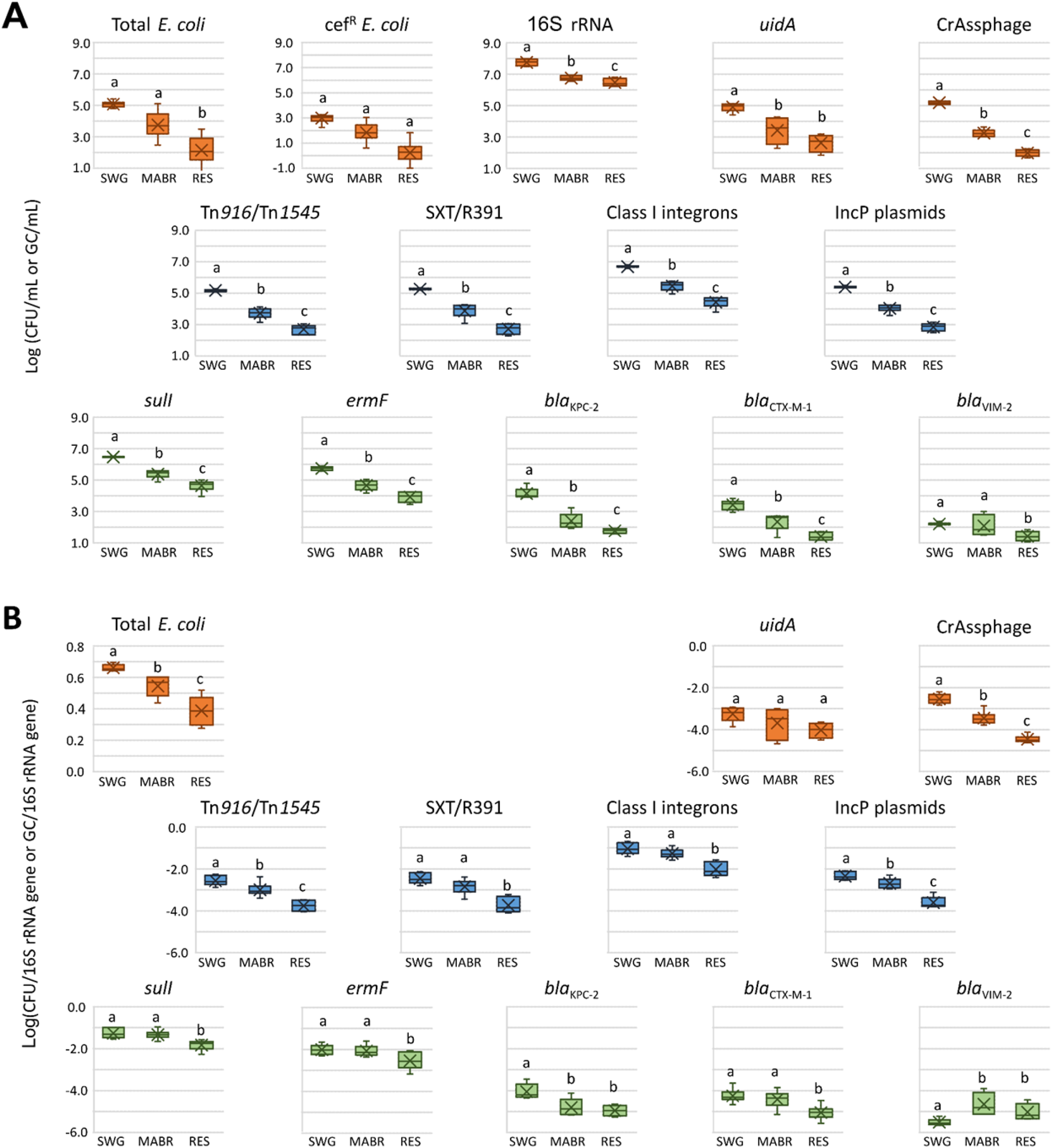
(A) Absolute and (B) relative abundances of total and cefotaxime-resistant *E. coli* and of 12 ARGs/MGEs markers. Each data point represents aggregated data collected from six different sampling times and four biological replicates. One-Way ANOVA Test followed by pairwise two sample t-test highlights significant differences. Colors represented different groups: MGE (blue), ARG (green) and bacterial (orange) markers and indicators (CrAssphage with CPQ_056 gene).

### ARG&MGE dynamics along the sewage-MABR-Reservoir continuum

The abundance of the nine targeted ARG&MGE markers was monitored in the raw sewage, MABR and reservoir by qPCR/ddPCR for six sampling dates (August 19^th^ and 25^th^; September 15^th^; November 24^th^; and December 1^st^ and 15^th^, 2020; **Figure. 3A**). In parallel, 16S rRNA, *uidA* and CrAssphage genes were monitored to estimate total bacterial, *E. coli* and *Bacteroides* phages, respectively.

The low standard deviations observed for most of the targeted genes indicate that the abundance of these markers is relatively steady over time in each compartment. On average, bacterial loads (estimated by 16S rRNA gene abundance) varied from 8.2 log-units ml^-1^ in sewage to 7.1 log-units ml^-1^ after MABR and 6.7 log-units in the reservoir, indicating that the reduction in bacterial abundance is higher in the MABR than in the reservoir. Apart from *bla_VIM-2_*, all the markers followed the same trend with an approximate 2-order of magnitude reduction across the continuum (**Figure. 3A** and **Figure. S5**). The normalized ARG&MGE abundance to 16S rRNA gene abundance (**Figure. 3B**) estimates the relative abundance of the different markers in the communities. With the exception of *bla*_VIM-2_, the relative reduction of all of the targeted genes and bacteria was higher in the reservoir than in the MABR. Furthermore, while the absolute abundance of SXT/R391, class 1 integrons, incP plasmids, *sul1, ermF* and *bla*_CTX-M-1_ decreased after MABR and reservoir, reduction in the relative abundance of these genes was only significant (*p*-value < 0.05) in the reservoir, thus suggesting a community shift.

### Microbial community composition along the sewage-WWTP-Reservoir continuum

#### Bacterial community composition

We evaluated bacterial community diversity (**Figure. S6**) and structure (**Figure. 4A**; **Figure. 4C**; **Figure. S7A**) in each of the three compartments. Average Shannon and Phylogenetic Diversity indices (**Table S4**) increased from 7.1 to 7.9, and 206 to 240, respectively, from the raw sewage to MABR effluents, but decreased in the reservoir. The phylum level evaluation of raw sewage, MABR and reservoir samples revealed distinct bacterial community profiles (**Figure. S7A**), but the reservoir microbiome displayed time-dependent variations that were substantially more dramatic. In the reservoir samples collected in November and December were richer in *Cyanobacteria* than in the previous sampling times. Family-level analysis suggested that *Cyanobacteria* were, indeed, predominantly chloroplasts (**Figure. S4A**), corresponding to eukaryotic algae also predominant in these microbial community profiles (see below). Members of the phyla *Pseudomonadota* (31.9% SWG, 37.7% MABR, 34.1% RES), *Bacteroidota* (11.0% SWG, 14.0% MABR, 15.4% RES) and *Actinobacteriota* (6.7% SWG, 10.9% MABR, 11.9% RES) were among the most represented in all samples, with the relative abundance of *Bacillota* and *Campylobacteriota* sharply decreasing from raw sewage to the reservoir (13.1% to 4.7% and 26.9% to 3.6%, respectively). In contrast, the relative abundance of other groups increased in the reservoir along the different sampling times. Most notably *Cyanobacteria* (ranging from 2.6% to 43.7%, in December), *Pseudomonadota* (ranging from 27.6% to 41.3 in August) and *Verrucomicrobiota* (ranging from 0.4% to 11.9 in September), in a pattern that was sampling-date-dependent.

**Figure 4.**
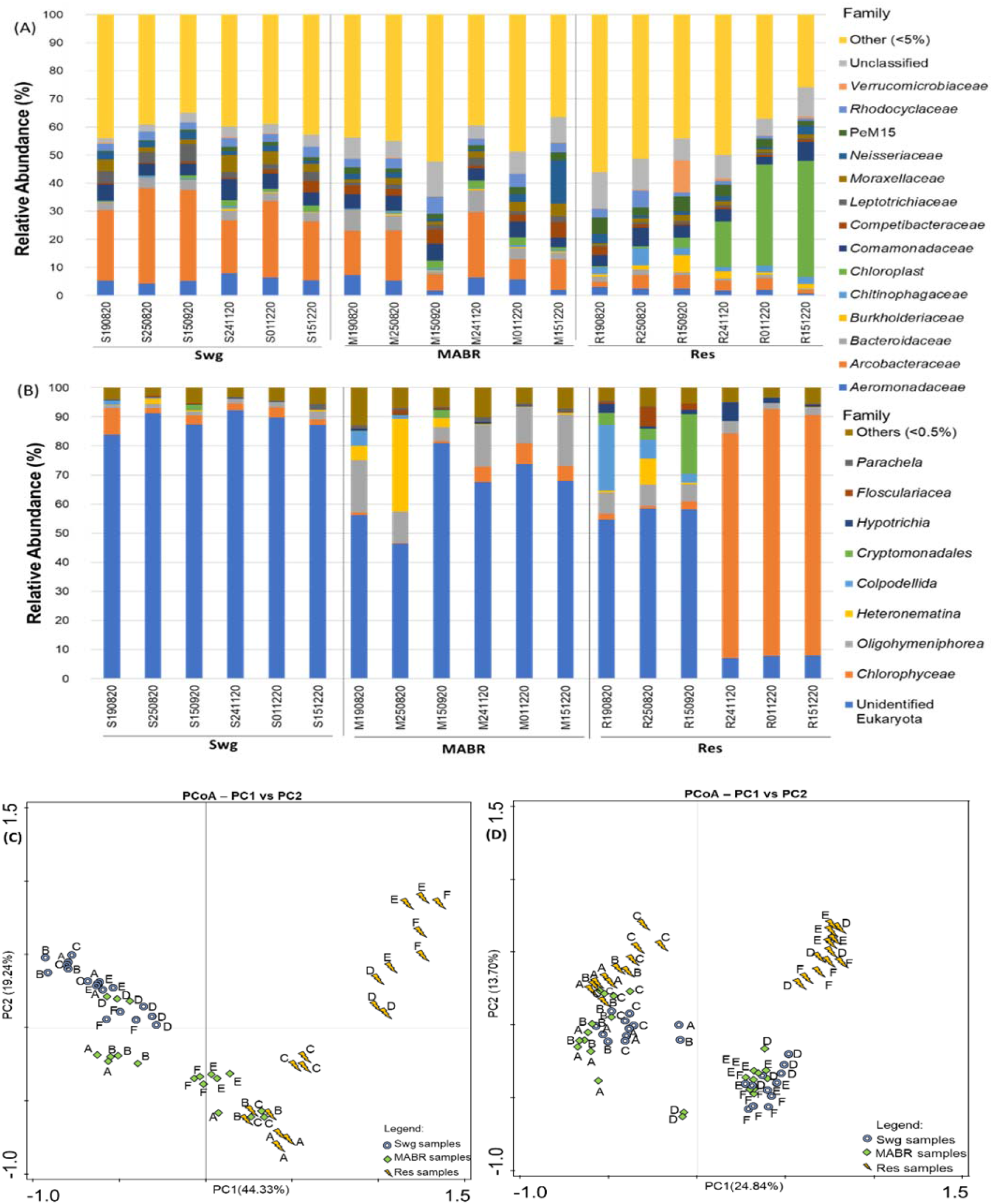
Bacterial and Eukarya community composition, structure and diversity. Family level taxonomy of Bacteria (A) and Eukarya (B), determined by targeting the V3-V4 variable regions of the 16S rRNA gene, and the V9 variable region of the 18S rRNA gene, respectively. Bars show taxa with relative abundance >5% and >1%, of defined bacterial and eukaryotic communities, respectively, in raw sewage (Swg), MABR or Reservoir (Res) samples. Prokaryotic (C) and Eukaryotic (D) diversity in raw sewage (Swg), MABR or Reservoir (Res) samples. Sampling dates from 2020 are identified from A to F: A – 19th August, B – 25th August, C – 15th September, D – 24th November, E – 1st December and F – 15th December

#### Eukaryotic microbial community composition

We evaluated the eukaryotic community composition in the raw sewage, MABR and reservoir, concomitant to bacterial community composition (**Figure. 4B**; **Figure. 4D**; **Figure. S7B**). Throughout the experiment the MABR was dominated by *Ciliophora* (13.4–25.8% rel. abundance), with the exception of a brief period when *Euglenazoa* (up to 32.9 ± 9.1% at 25^th^ August 2020) and *Ochrophyta* (up to 62.8 ± 9.3% at 15^th^ September 2020) became dominant (**Figure. 4B**), corresponding to the above described MABR malfunction. The reservoir was initially dominated by the bacterivorous protists *Ciliophora* (38.8 ± 2.0%) and *Proteoalveolata* (22.5 ± 2.3%). As indicated above, *Ciliophora* was the dominant eukaryote in the MABR, and it is possible that they initially colonized the reservoir. However, the relative abundance of both groups significantly decreased with time (p=0.02; n=24; non-parametric test for association based on Spearman’s rho), with *Proteoalveolata* dropping below detection levels, and the relative abundance of *Ciliophora* dropping to 5.2 ± 0.6 %. Conversely, the relative abundance of *Chlorophyta* significantly increased over time (p<0.001; n=24; ANOVA) and became the dominant phyla from November onward, accounting for more than 75% of the relative abundance of eukaryotes in the reservoir, strongly correlating to the increase in relative abundance of chloroplasts described above. This increase in *Chlorophyta* abundance resulted in a significant shift in eukaryotic community composition (p<0.01; n=24; AMOVA (Analysis of MOlecular VAriance) of NMDS distances; **Figure. 4B**) relative to the three early profiles where blooms of this group of green algae were not observed in the reservoir. In contrast to the reservoir, no significant temporal differences were observed in the raw sewage and MABR eukaryotic communities, supporting the trend reported above in which microbial community dynamics in the reservoir were independent of those in the MABR or the sewage.

#### Assessing correlations between microbial, physiochemical and environmental parameters

Redundancy analysis (RDA) was conducted to investigate possible statistically significant correlations (p< 0.01) between physicochemical parameters, ARGs and MGEs, and the microbial community composition in the raw sewage, MABR and reservoir (**Figure. 5**, **Figure. S8**, **Figure. S9**). Since bacterial and ARG and MGE marker removal in the MABR was at least partially associated with biomass removal, we focused on trends that occurred between the MABR and the reservoir. While correlation does not necessarily indicate causation, we observed trends that potentially shed light on the complex abiotic and biotic processes that occur in the two modules and how they affect the system dynamics. The reduction of total bacteria observed in the reservoir (as measured by 16S rRNA gene abundance), total coliforms, total *Escherichia coli* (including *uid*A), and all the measured ARGs (except *bla*_VIM-2_) and MGEs was positively correlated (p< 0.01) to pH, and negatively correlated to electric conductivity, total nitrogen and dissolved oxygen. The photosynthetic eukaryotic taxa *Dinoflagellata, Proteoalveolata* and *Ochrophyta* strongly correlated (p< 0.01) to the reservoir, as did the bacterial phylum *Actinobacteriota*. In contrast, the non-photosynthetic *Euglenozoa* were significantly associated (p< 0.01) with the MABR.

**Figure 5.**
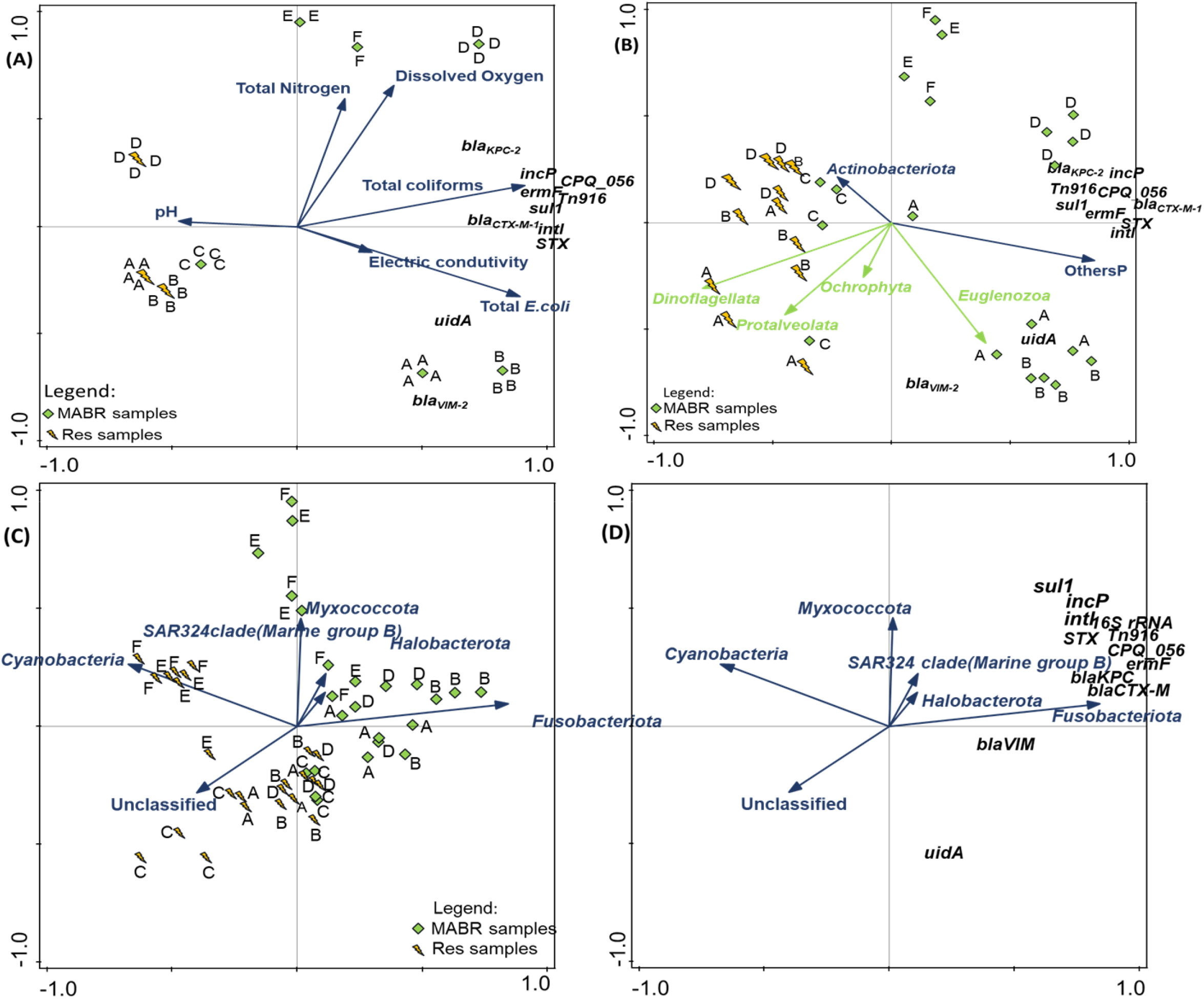
Redundancy analysis (RDA) of the variation of ARGs, MGEs and prokaryotic and eukaryotic populations in the MABR and reservoir samples. Sampling dates from A to F: A – 19^th^ August, B – 25^th^ August, C – 15^th^ September, D – 24^th^ November, E – 1^st^ December and F – 15^th^ December. (**A**) – RDA of the variation ARGs, MGEs and *uidA* gene in MABR (M) and reservoir (R) samples in function of measured physico-chemical parameters. The test variables (ARGs, MGEs and *uidA)*are represented in black and the explanatory variables in blue (physico-chemical parameters). (**B**) - RDA of the variation ARGs, MGEs and *uidA* gene in function of the prokaryotic and eukaryotic community phyla members with relative abundance >5% and >1%, respectively, and summed as others (E, eukaryote or P, prokaryote) for lower values. Additional information and statistical analyses are provided in **Table S9**. (C) and (D) - Redundancy analysis (RDA) of the variation ARGs, MGEs and 16S rRNA genes in M and R samples in function of prokaryotic community. The test variables (ARGs, MGEs and uidA) are represented in black and the explanatory (prokaryotes) in blue. The explanatory variables were associated with 78.7% of the observed variation among the test variables.

## Discussion

This study unveiled three major aspects of bacterial and ARGs/MGEs dynamics that are discussed below – i) “Real life” pilot systems provide novel insights into antibiotic resistance dynamics that are not obtainable through analysis of full- or bench-scale systems; ii) Biomass removal is pivotal for the extensive elimination of ARB and ARGs and MGEs in secondary wastewater treatment; iii) polishing of treated wastewater, to remove additional ARB, ARGs and MGE loads, can be facilitated by exploiting ecosystem services provided by open water storage reservoirs. Each of these ae discussed below.

### “Real life” pilot systems as essential tools for wastewater ecology studies

Raw sewage and secondary effluents constitute a source of fecal-derived ARB and associated ARG/MGE contamination that can be released to the environment and contribute to the global scope of antibiotic resistance. Nonetheless, there is a lack of holistic understanding regarding the dynamics and fate of ARB and ARGs in receiving aquatic ecosystems due to the complexity of receiving ecosystems, which does not enable differentiation between effluent discharge and other anthropogenic sources. Bench-scale experiments shed-light on specific factors that influence antibiotic resistance determinants in WWTPs and receiving environments (31, 32), but they may not accurately predict large-scale processes, especially when synthetic wastewater is used instead of real sewage. While the reservoir is not completely representative of large-scale effluent reservoirs or natural limnological ecosystems, its size, controlled and contained structure, and influx of real secondary effluents, provided insights unobtainable with either bench- or full-scale systems. This was augmented by the inclusive methodological approach applied that coupled evaluation of total and cefotaxime-resistant coliforms and *E. coli*, quantification of rank I to IV ARGs as well as a panel of MGEs, bacterial and eukaryotic community composition and a suite of physicochemical parameters, all investigated in multiple profiles over a six-month period. We recommend that studies aiming to understand the dynamics of ARB and ARGs in effluent receiving environments adopt the use of that large-scale pilot reservoirs operating in the field.

### Mitigation of ARB and ARGs in the MABR is primarily attributed to biomass removal

Raw sewage entering the MABR was highly uniform, microaerophilic and contained high levels of total nitrogen (primarily ammonia) and organic carbon. Aeration in the MABR increased oxygen levels, facilitating a sharp reduction in dissolved organic matter (and turbidity) and total nitrogen, which was attributed to gasification of N_2_ and CO_2_ through combined nitrification/denitrification on the MABR membranes. The total and cefotaxime-resistant *E. coli* and ARG and MGE loads in the MABR dropped by approximately 1.5 log units ml^-1^ between sewage and MABR, but the relative abundance (normalized to 16S rRNA gene levels) of most ARG and MGE markers, and *E. coli* were not significantly reduced. The bacterial community composition of the MABR effluent was much closer to that of the raw sewage than to the MABR. This is supportive of previous studies (33–36) reporting that microbiomes of secondary WWTP effluent are significantly different from activated sludge, and actually more closely resemble influent microbiomes, suggesting that while ARB and ARGs are removed in the secondary treatment process, their general composition is similar to that of the sewage. The conclusions of this study suggest that in future studies, secondary treatment biosolids should be analyzed along with effluents in order to perform a complete mass balance, and determine how much nitrogen and carbon were gasified *vs*. how much remained in the sludge, or to evaluate the microbial community composition of the sludge relative to the aqueous effluent.

### Reservoir dynamics suggests natural attenuation facilitated by a resilient microbiome

Abundance of ARG and MGE markers and *E. coli* further decreased in the reservoir, but in contrast to the MABR, for the most part, the relative abundance (normalized to 16S rRNA genes) also dropped significantly. The reservoir microbiome was significantly different to the MABR microbiome, with higher relative abundance of *Pseudomonadota* and *Acidobacteriota*. The coupling of 16S and 18S rRNA gene analyses revealed that the sharp increase in *Cyanobacteria* observed in December 2020 was, in essence, chloroplast, predominantly of the green algal family *Chlorophyceae*. It is unclear whether proliferation of these green “blooms” was linked to reduced temperatures in December, to the fact that it took time for these populations to mature, or to other unknown stochastic or deterministic causes. Similar algae blooms were also observed in a previous study that evaluated plankton community changes in a freshwater reservoir amended with treated wastewater effluents (Teltsch et al. 1992), and thus future studies need to identify specific parameters that induce their proliferation, and determine why cyanobacteria blooms were not encountered in the reservoir.

The removal of ARB and ARG markers in the reservoir was supported by a previous study that observed similar trends in a commercial-scale effluent reservoir used for irrigation (11). We attribute this mitigation to ecosystem resilience (37), attributed at least in part to natural attenuation facilitated by a highly resilient “environmental microbiome” that outcompete and prevent colonization of effluent-derived ARB and associated ARGs as previously described (38). Chen *et al*. showed that reduction of microbial diversity exacerbates the spread of antibiotic resistance in soil (39), highlighting its importance to resilience and resistance of receiving environments towards fecal derived ARB and ARGs. Interestingly, relative reduction in *E. coli* and ARG and MGE levels was relatively stable in the reservoir, despite strong temporal fluctuations in microbial community composition, suggesting functional redundancy of native environmental communities (40). These “native” communities undoubtedly play a role in competitive exclusion of sewage-derived ARG-harboring bacteria, which is most likely facilitated by a myriad of mechanisms including adaptation to abiotic stressors (*i.e*. radiation, temperature), resource competition and antibiosis. Amplicon sequencing only provides relative abundance values and future studies need to go beyond the community level and into the functional realm in order to pinpoint genes associated with specific characteristics (*i.e*. metabolic pathways, stress related proteins, antibiotics etc.) that facilitate the observed ecological resilience. Furthermore, batch experiments using synthetic communities should be conducted to assess the capacity of specific population/communities to mitigate specific ARB and ARGs, and these should be complemented with relevant Eukaryotic grazers to evaluate the role of predation.

Contrary to the other ARGs, in some of the analyzed profiles the relative abundance of *bla*_VIM-2_ seemed to increase in the MABR and did not significantly decrease in the reservoir. The temporal dynamics of this gene revealed a large standard deviation corresponding to a temporal shift from higher *bla*_VIM-2_ abundance in MABR effluent *vs*. raw sewage in August 2020, to an expected removal/depletion profile from September on. While the specific factors responsible for this phenomenon are unknown, it may be suggested that in the analyzed system *bla*_VIM-2_ is primarily harbored by bacteria that colonize the MABR that were potentially washed out in the early phase of the experiment (explaining their abundance in the reservoir). The fact that *bla*_VIM-2_ was not significantly removed in the reservoir, suggests that the bacteria that harbor it are much more resilient in aquatic ecosystems than bacteria harboring other “hazardous”β-lactamases such as *bla*_KPC-2_ and *bla*_CTX-M-1_. Although ubiquitous to several Gram-negative taxa, *bla*_VIM-2_ was first isolated (41) and is often detected in wastewater-associated and environmental biofilm-producing *Pseudomonas* spp. strains (42). The unorthodox behavior of *bla*_VIM-2_ underlines the fact that ARGs need to be monitored individually when conducting source tracking, and that the relation between individual ARGs and specific phyla and MGEs need to be better understood to enable accurate risk assessment.

### Effluent storage reservoirs as a means of mitigating antibiotic resistance

Our results highlight the robust ARB and ARG removal capacity of effluent storage reservoirs, corroborate a previous study which reported that stabilization reservoirs used for effluent storage prior to irrigation removed fecal coliforms by up to five orders of magnitude before chlorination (43), depending on retention time and operational conditions. Removal rates of ARGs by the reservoir are comparable to more sophisticated disinfection processes such as chlorination (44) and UV (45), suggesting that they may provide a simple and “environmentally sustainable” alternative. The capacity of autochthonous microbiota to hamper the proliferation of invasive bacteria has been increasingly demonstrated and debated in the scientific literature and should be a major research focus on antibiotic resistance combat in the environment (46, 47). Overgrowth control of potentially hazardous bacteria during storage of ozone treated wastewater through natural competition. Although primarily developed for storage of recycled effluents used for irrigation, our results suggest that if properly configured, reservoirs may also be applied to polish effluents prior to discharge into aquatic ecosystems.

## Conclusions

Temporal analysis revealed that both membrane-based secondary municipal sewage treatment and subsequent reservoir storage mitigated antibiotic resistant bacteria and associated ARGs and MGEs. However, in the MABR the phenomenon was primarily facilitated by biomass removal, whereas in the reservoir it was attributed to shifts in microbial community composition towards a more resilient “environmental” microbiome that seemingly prevents colonization of fecal derived bacteria and associated genes through competitive exclusion associated with ecosystem resilience. Based on these results, we do not only propose the implementation of reservoirs for storing effluents used for irrigation, but also as a means of mitigating fecal bacteria and antibiotic resistance in effluents discharged to aquatic environments, especially in developing regions that lack sophisticated wastewater treatment infrastructure.

## Acknowledgments

This study was supported by the European Union’s Horizon 2020 research and innovation programme project “DSWAP” under the PRIMA program under grant agreement No 1822. CM, UK & TUB were also supported through the ANTIVERSA project funded by the Agence Nationale de la Recherche (France), and the Bundesministerium für Bildung, und Forschung (Germany), respectively, under grant number 01LC1904A. CM received additional support from the LTSER-France, and the Lorraine Region through the research network of Zone Atelier Moselle (ZAM). Responsibility for the information and views expressed in the manuscript lies entirely with the author(s).

